# The speed limits for tau pathology progression in Alzheimer’s disease

**DOI:** 10.1101/2024.09.02.610787

**Authors:** Merle C. Hoenig, Verena Dzialas, Elena Doering, Gérard N. Bischof, Thilo van Eimeren, Alexander Drzezga, Alzheimer’s Disease Neuroimaging Initiative

**Affiliations:** Research Center Juelich, Institute for Neuroscience and Medicine II, Molecular Organization of the Brain, Juelich, Germany; University of Cologne, Faculty of Medicine and University Hospital Cologne, Department of Nuclear Medicine, Cologne, Germany; University of Cologne, Faculty of Mathematics and Natural Sciences, Cologne, Germany; University of Cologne, Faculty of Medicine and University Hospital Cologne, Department of Neurology, Cologne, Germany; German Center for Neurodegenerative Diseases, Bonn/Cologne, Germany

## Abstract

**Objective:** To examine interactive effects of modifiable factors, genetic determinants and load-dependent pathology effects on tau pathology progression.

**Methods:** Data of 162 amyloid-positive individuals were included, for whom longitudinal [18F]AV-1451-PET scans, baseline information on global amyloid load, ApoE4 status, body-mass-index (BMI), hypertension, education, neuropsychiatric symptom severity and demographic information were available in ADNI. All [18F]AV-1451 PETs were intensity-standardized (reference: inferior cerebellum), z-transformed (control sample: 147 amyloid-negative subjects) and subsequently thresholded (z-score > 1.96) and converted to volume-maps. Based on these volume-maps, tau-changes over time were assessed in terms of 1) *tau-speed* (i.e. newly affected volume at follow-up), and 2) *tau-level-rise* (i.e. tau increase in previously affected volume). These two measures were entered as dependent variables in separate linear mixed effects models including four baseline risk factors (BMI, education, hypertension, neuropsychiatric symptom severity), baseline amyloid, tau-volume or tau burden, ApoE4 status, clinical stage, sex, and age as predictors. Next, we tested the interactive effects between baseline amyloid or tau burden with the four modifiable factors on either tau-speed or tau-level-rise, respectively.

**Results:** Faster tau-speed was linked to higher BMI, female sex, ApoE4-status, and baseline tau-volume. The effect of baseline tau-volume on tau-speed was driven by greater global amyloid burden. In terms of tau-level-rise, we observed that lower hypertension and BMI were linked to a slower increase in tau burden. A load-dependent effect of baseline amyloid and tau burden was found. Higher amyloid and BMI as well as lower education and higher tau burden were linked to greater tau-level-rise.

**Conclusion:** Education, BMI and hypertension differentially influence tau speed and level rise by its interaction with initial pathological burden. Timely modification of these factors may overall slow tau’s progression.

## Introduction

The recent proposition by the prevention commission of Livingston and colleagues suggests that nearly 50 percent of all dementia cases could be prevented by modifying 14 risk factors ^1^. The report suggests that, for instance, ensuring good quality of education, treating depression effectively, preventing hypertension, and maintaining a healthy weight in midlife could effectively reduce the overall dementia risk. This suggests that these factors are inevitably linked to the onset of Alzheimer’s disease (AD) and may thus also potentially prolong the progression of AD pathology.

The breakthrough in *in vivo* imaging of the neuropathological hallmarks of AD, namely amyloid and tau pathology with PET imaging, has revealed that symptomatic AD progression is more closely linked to tau than amyloid burden ^2^. Yet, amyloid-ß load (globally^3,4^ or in selected regions^5^) remains the strongest predictor of tau aggregation and spreading, compared to genetic factors (e.g., ApoE4) and other vulnerabilities like age or white matter hyperintensities ^6,7^. However, little is known about how modifiable risk factors, such as education, depression, body-mass-index (BMI) or hypertension, interact with baseline pathology burden and subsequent tau spreading. Cross-sectional studies have so far suggested that higher education ^8,9^, neuropsychiatric symptoms ^10^, higher BMI ^11^, and hypertension^12^ are associated with greater AD pathology. Yet, in terms of the effects of late-life BMI, findings have been more inconsistent ^13,14^.

In this study, we focus on examining the concomitant influence of four potentially modifiable risk factors (education, neuropsychiatric symptom severity, BMI, hypertension) to tau pathology spreading over time. This volumetric approach has recently been shown to be more closely linked to disease progression than intensity measures^15,16^. Specifically, we tested the contribution of these factors to the spatial progression of tau pathology (i.e. tau speed) and the local increase in tau burden (i.e. tau level rise). We anticipated that higher education, lower BMI, an absence of hypertension and neuropsychiatric symptoms would be associated with slower tau speed and lower levels of tau level rise, while baseline amyloid and tau burden, and ApoE4 carriership (i.e., established contributors) would be inversely related with tau speed and level rise. Additionally, we assumed that the protective effects of these modifiable factors would be driven by an interaction with lower baseline amyloid burden, rather than baseline tau burden, presuming that amyloid burden is they key driver of tau spreading.

## Materials & Methods

### Participants

Data for this study were retrieved from the Alzheimer’s Disease Neuroimaging Initiative (ADNI, adni.loni.usc.edu) in July 2024. Key inclusion criteria were: 1) at least two evaluable [18F]AV-1451 PET scans, 2) baseline amyloid PET scan within three months of the baseline tau PET acquisition & confirmed amyloid positivity, 3) ApoE4 carriership information, 4) available baseline information on BMI, hypertension, education and neuropsychiatric inventory (NPI) in addition to demographic information. This resulted in 162 individuals fulfilling the criteria in ADNI: 77 cognitively-unimpaired (CU), 55 patients with mild cognitive impairment (MCI) ^17^, and 30 patients with AD ^18^. Demographic characteristics of the respective groups are provided in Table 1. In addition, an amyloid-negative cognitively normal group was included, for which two tau-PET scans and a baseline amyloid scan was available *(N=147; M(age)=71*.*63±6*.*32, M(education)=16*.*92±2*.*38, M(ADAS13)=7*.*56±4*.*28, M/F=67/80, ApoE4 (Yes, No, Missing)=45/100/2)*. This group was used as reference group for the z-transformation of the tau PET scans. Ethics committee approval was obtained at each of the participating centers of the Alzheimer’s Disease Neuroimaging Initiative (ADNI).

**Table 1.**
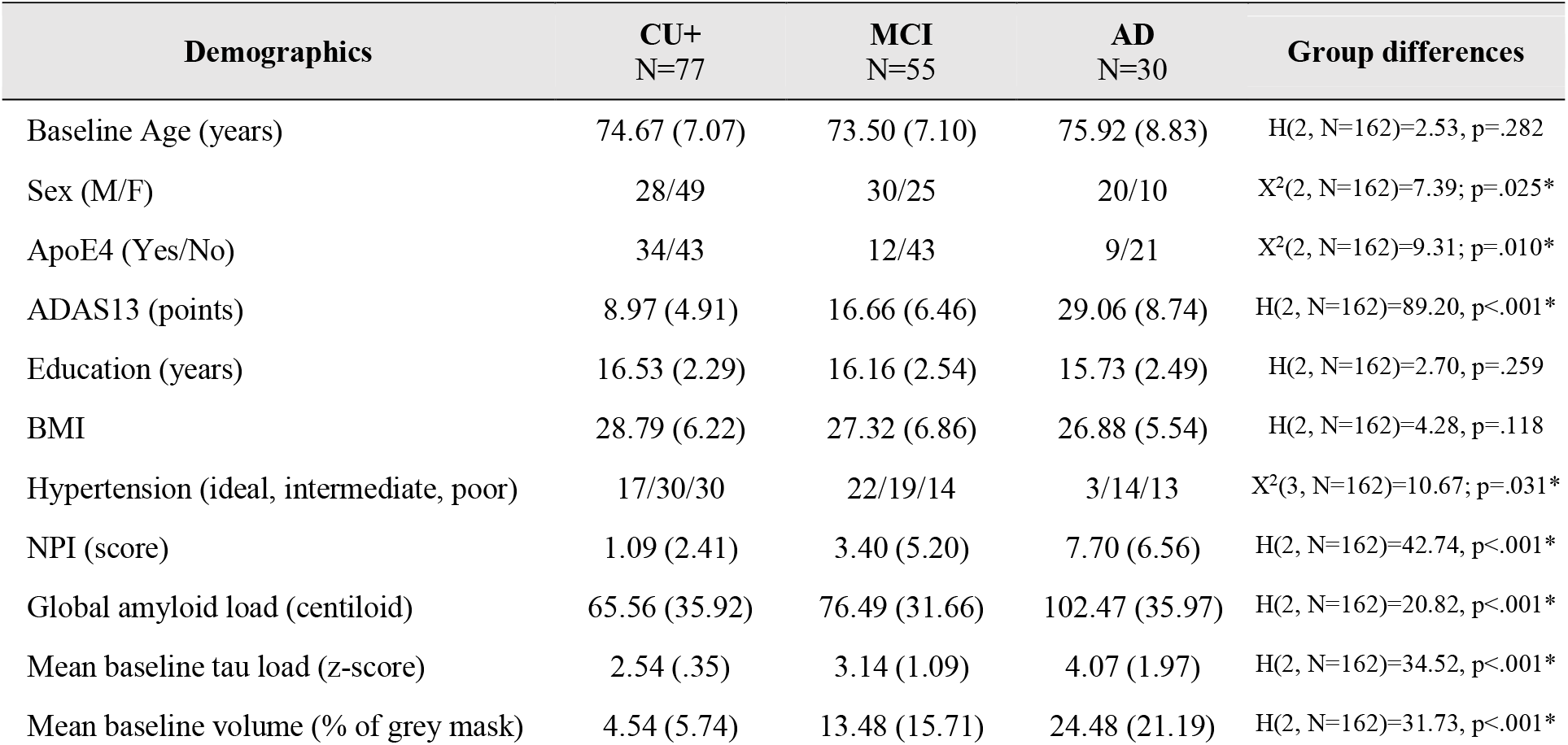
Demographic characteristics of the amyloid-positive sample. Mean and standard deviations are provided for continuous and distributions for ordinal variables. Significant group differences were tested using a Kruskal-Wallis-test for continuous and Chi-squared test for ordinal variables.

#### Risk factors

##### Education

Educational attainment comprised years of formal school education in addition to following higher education.

##### BMI

Height (inches or centimeters) and weight (pounds or kilograms) were converted to meters and kilograms, respectively. BMI was calculated: weight (kg)/ height (meters)^2^.

##### Hypertension

Hypertension information was derived based on the systolic (SBP) and diastolic (DBP) blood pressure assessed at the respective baseline visit. Following groups were established using the cut-off values based on the recommendations of the American Heart Association: ideal=<120 mmHg/<80 mmHg; intermediate= SBP 120-139 mmHg or DBP 80-89 mmHg; poor= SBP >= 140 mmHg or DBP >= 90 mmHg.

##### Neuropsychiatric inventory (NPI)

The NPI entails ten behavioral areas and two neurovegetative areas. Symptoms need to be present for at least four weeks and scoring is based on *frequency* ranging from 1-4 and *severity* ranging from 1-3. A total NPI score is then calculated by adding the scores (*frequency by severity)* of the twelve domains, which was used in this study to approximate neuropsychiatric symptom severity.

#### PET image processing

Pre-processed tau PET images provided by ADNI were used, including co-registration to the individual MRI, normalization to MNI space and intensity-normalization using the inferior cerebellar grey. The respective tau PET scans (baseline and FUs) were z-transformed using the mean of the baseline and first FU tau PET scans of the amyloid negative control group to control for repeated measurement effects. Information on global amyloid load at baseline was extracted based on the centiloid information provided in the data sheet “UCBERKELEY_AMY_ 6MM”. We used centiloids as a tracer-independent measure for global amyloid as either [18F]Florbetapir or [18F]Florbetaben PET scans were available.

#### Volumetric approach

We performed a volumetric approach to quantify the speed (spatial extent) and level rise (increase) of tau pathology over time. First, we thresholded the tau z-maps at a z-score of z >= 1.96 (one-tailed p-value < .025) and subsequently created binarized volume maps comprising only voxels above this threshold. Then, we defined two respective volumes for each follow-up time point (FU_tx_):

1. *Newly affected volume* = Binarized z-map FU_tx_ – binarized z-map of baseline tau scan; FU_Volume_tx_ -BL_Volume_. Consequently, the newly affected volume reflects all regions that were not affected at baseline, but only at FU. Number of voxels were then translated into volume in liters (voxel number * (voxel size of 1.5mm)^3^/1,000,000). This measure was used as proxy for the *speed* of tau pathology over time.
2. *More affected volume* = Regions of spatial overlap at FU_tx_ with previous time point; FU_Volume_tx_ ∩ FU_Volume_tx-1_. This volume was used to quantify the increase in tau pathology over time in regions that had already been affected at the previous time point. The z-score change in the more affected volume was extracted by taking the mean difference of FU_tx_ and FU_Baseline_ within the more affected volume between the two time points. The z-score changes was used to quantify *tau level rise*.

Additionally, the baseline volume and mean z-score were extracted for each individual.

#### Statistical Analyses

##### Correlation analyses and group-comparisons to test baseline associations

To investigate the relationships among variables of interest - BMI, educational attainment, NPI score, baseline centiloid value, and mean baseline tau z-score - we used partial Spearman correlations (one-tailed), adjusted for age and sex. For categorical variables such as hypertension and ApoE4 carriership, we employed the Kruskal-Wallis test, with centiloid value, baseline tau burden, volume, BMI, education, and NPI as the dependent variables. Chi-squared tests were used to compare the distribution of ApoE4 and hypertension severity. Correction for multiple comparisons was performed using Benjamini-Hochberg correction (false discovery rate).

##### Linear mixed effects modeling (LMM)

To examine the effects of the (non-)modifiable factors to the speed and level rise of tau pathology, we performed linear mixed effects modeling in SPSS 28 (IBM Corp., Armonk, NY USA).

The first LMM model included the newly affected volume (i.e. tau speed) in up to five year follow-up data (**Model 1 - Tau Speed**). The following continuous fixed effects were introduced: age at baseline, time difference in months from baseline tau scan, baseline centiloid, baseline tau volume, education years, BMI and NPI total score. Ordinal fixed effects comprised sex (F/M), ApoE4 carriership (yes/no), group status (CU, MCI, AD), hypertension (ideal, intermediate, poor). In addition, the model included two random effects (subject and time) that allowed individual intercepts and slopes using an unstructured covariance matrix. Given our a priori hypotheses, we subsequently added the following interaction terms to the model: baseline centiloid and baseline tau volume with the four risk factors, respectively in addition to the interaction between hypertension severity and ApoE4 status given previously reported interaction between these two variables on amyloid burden ^19^.

The second LMM included the z-score change over time in the more affected volume as dependent variable (**Model 2 - Tau Level Rise**). The fixed effects remained the same as in Model 1, except that we included baseline tau burden instead of baseline tau volume as fixed effect since we focused on tau intensity in this model. Given subjects did not significantly vary in their intercept values (tau burden at baseline), we only introduced a random slope. Next, we tested the same interaction effects as in model 1, except that we used tau burden instead of tau volume in the interactions.

As we tested the effects of baseline tau volume or tau burden on the subsequent increase in the newly affected volume or tau increase in the more affected volume, for some individuals (n=72) only one FU time point was available. As we wanted to rule out a potential effect of these singular measurements on the LMMs, we ran the same analyses with individuals (n=90) that had at least two FUs available.

Normal distribution for models’ residuals was tested. Significance level was set at p < .05. Results are reported as unstandardized beta-coefficients (ß), p-values and corresponding confidence intervals (CI). Visualization of results was performed in R Studio using *ggplot* ^20^.

## Results

### Group characteristics

The three groups (CU, MCI, AD) of interest did not differ significantly in terms of age, education and BMI, but in the remaining variables (sex, ApoE4 carriership, hypertension, ADAS13, centiloid value, baseline tau burden and volume). The mean ranks per group were related to AD severity (CU<MCI<AD). There were more ApoE4 carriers in the MCI group compared to the CU and AD group and more men in the MCI and AD group in comparison to the CU group. Hypertension (intermediate and poor) showed a higher frequency across groups in comparison to ideal hypertension (see table 1).

### Baseline associations of risk factors with AD pathology

Significant correlation effects were observed between baseline amyloid burden and tau burden (rho=.459, p<.001), tau volume (rho=.346, p<.001), and the NPI (rho=.219, p=.003), whereas amyloid burden was negatively associated with BMI (rho=-.157, p=.024) and education (rho=-.150, p=.029). Baseline tau burden was positively correlated with tau volume (rho=.787, p<.001) and NPI (rho=.209, p=.004). Likewise, baseline volume was positively associated with the NPI (rho=.208, p=.004). BMI was significantly correlated with NPI (rho=-.137, p=.042). Remaining correlations did not yield significance. After correction for multiple comparison, the significant associations between amyloid burden, tau burden, tau volume as well as their associations with the NPI remained. The association of BMI (q= .051) or education (q= .054) with amyloid burden only showed a trend significance.

Greater hypertension severity was significantly associated with lower education (H(2; n=162)=6.688, p=.035), but did not survive FDR correction. ApoE4 carriership was related with baseline tau volume (H(1; n=162)=4.957, p=.026) and trend significantly linked with amyloid burden (H(1; n=162)=3.628, p=.057), which both did not survive multiple comparison correction. Results of all tested associations can be found in the supplementary information in S-Table 1.

### Tau Speed – Its de- and accelerating factors

Model 1 yielded a significant main effect of ApoE4 carriership (p=.039), BMI (p=.021), baseline tau volume (p<.001) and time (p<.001) on the newly affected volume over time. More specifically, ApoE4 carriership was linked to a greater tau speed. Likewise, BMI, tau volume and time were positively associated with the newly affected volume. The remaining variables were not significant (see Table 3). The interaction analyses revealed a significant effect of baseline amyloid and tau volume (β=-.005, p<.001, 95% CI, -0.008- -0.002) on the newly affected volume. Closer inspection yielded that greater centiloid levels and tau volumes at baseline were linked to the strongest increase in the newly affected volume. The interaction between centiloid levels and BMI resulted in a trend significant finding (β=.0001, p=.086, 95% CI, -0.0001-0.0001), while the remaining tested interaction terms were not significant. The LMMs in the subcohort with at least two FU time points yielded the same results, except that sex turned out to be a significant fixed effect with females presenting a greater increase than men (S-Table 2 & 3).

**Table 3.**
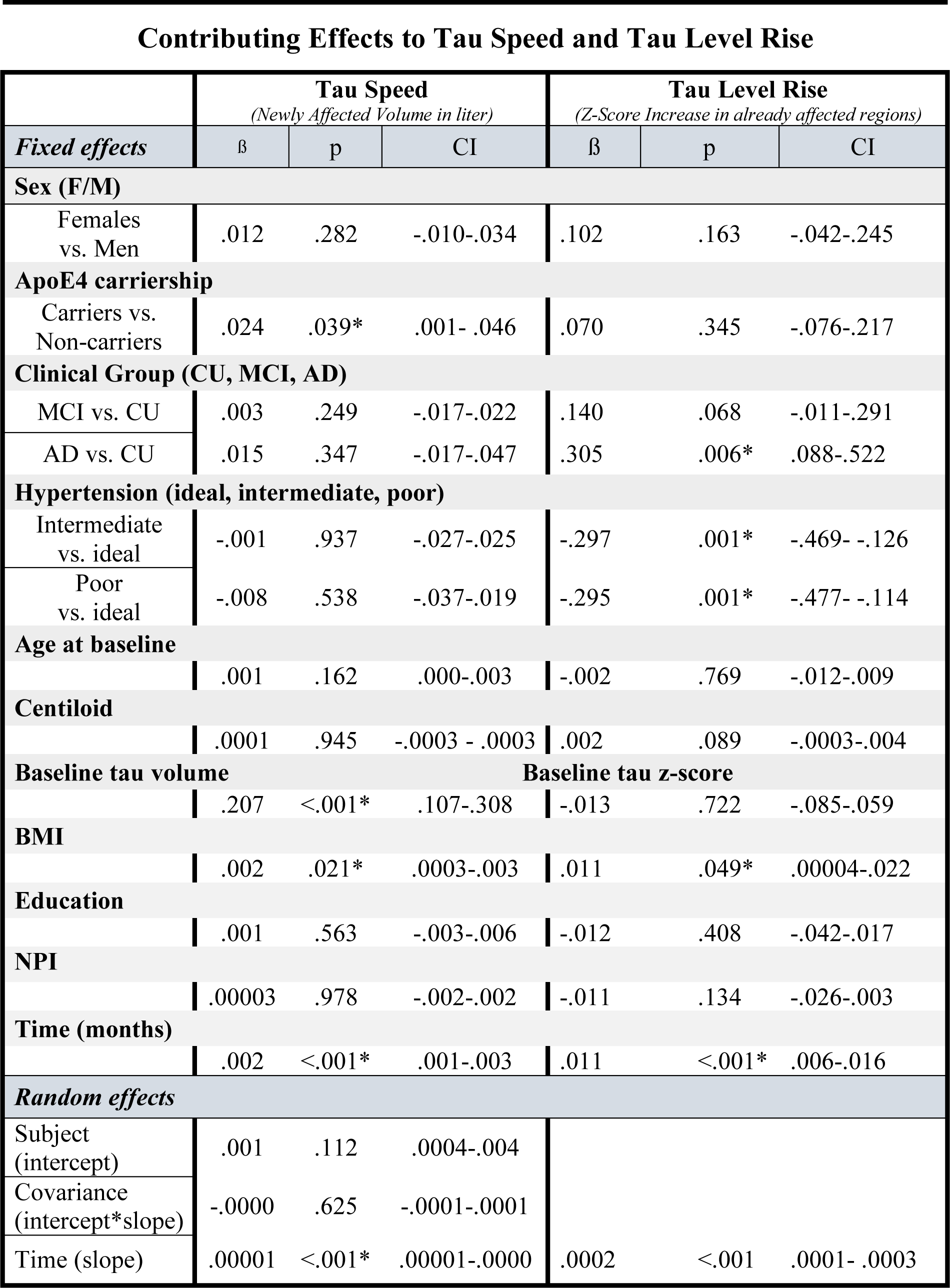
Results of the LMM for the newly affected volume and the z-score increase in the more affected volume over time. Significant fixed effects are highlighted with an asterisk. The effects of the random slopes are additionally provided for each model. Unstandardized beta-coefficients, p-values and confidence intervals (95%) are provided for all effects. The results relate to the cohort of 162 subjects.

**Table 4.** Interaction effects of the respective models. The fixed effects were the same as in Table 3, but the interaction terms were additionally entered into the respective models. Significant associations are highlighted with an asterisk. The F-statistic and p-values are provided for all interaction effects.

| *Interaction effects* |  | F | p |
| --- | --- | --- | --- |
| ***Model 1 - Tau Speed (Newly Affected Volume)*** |  |  |  |
| Centiloid | Tau volume | 14.46 | <.001* |
|  | BMI | 3.08 | .082 |
|  | Education | .806 | .371 |
|  | NPI | .187 | .666 |
|  | Hypertension | .215 | .807 |
| Tau Volume | BMI | .600 | .440 |
|  | Education | .806 | .371 |
|  | NPI | .559 | .459 |
|  | Hypertension | .785 | .458 |
| Hypertension | ApoE4 | .930 | .397 |
| ***Model 2 - Tau Level Rise (Z-Score Change in More Affected Volume)*** |  |  |  |
| Centiloid | Tau volume | 9.10 | .003* |
|  | BMI | 14.90 | <.001* |
|  | Education | .396 | .530 |
|  | NPI | .089 | .765 |
|  | Hypertension | .139 | .870 |
| Tau Burden | BMI | 2.432 | .120 |
|  | Education | 5.47 | .020* |
|  | NPI | .044 | .833 |
|  | Hypertension | .232 | .703 |
| Hypertension | ApoE4 | 2.37 | .095 |

### Tau Level Rise – Its de- and accelerating contributors

The tau level rise model yielded a significant effect of dementia group (p=.020), hypertension (p=.001), BMI (p=.049) and time (p<.001). The AD group presented a greater tau level rise over time in comparison to the CU group (p=.006). A trend significance for the comparison MCI against CU was found (p=.068). Unexpectedly, the poor (p=.001) and intermediate hypertension (p=.001) group presented a generally lower increase in tau pathology over time in comparison to the ideal hypertension group. Nonetheless, the poor hypertension group presented the steepest slope upon visual inspection. The remaining variables did not reach significance. Significant interaction effects were observed for: Baseline centiloid and tau burden (p=.003), baseline centiloid and BMI (p<.001), and baseline tau burden and education (p=.020). The remaining interactions did not yield significant results. Again, results could be replicated in the smaller subcohort (S-Table 2 & 3).

A summary of the LMM’s model statistics can be found in Table 2 and 3 for tau speed and tau level rise. Significant effects are summarized in Figure 1 and visualized in Figure 2.

**Figure 1.**
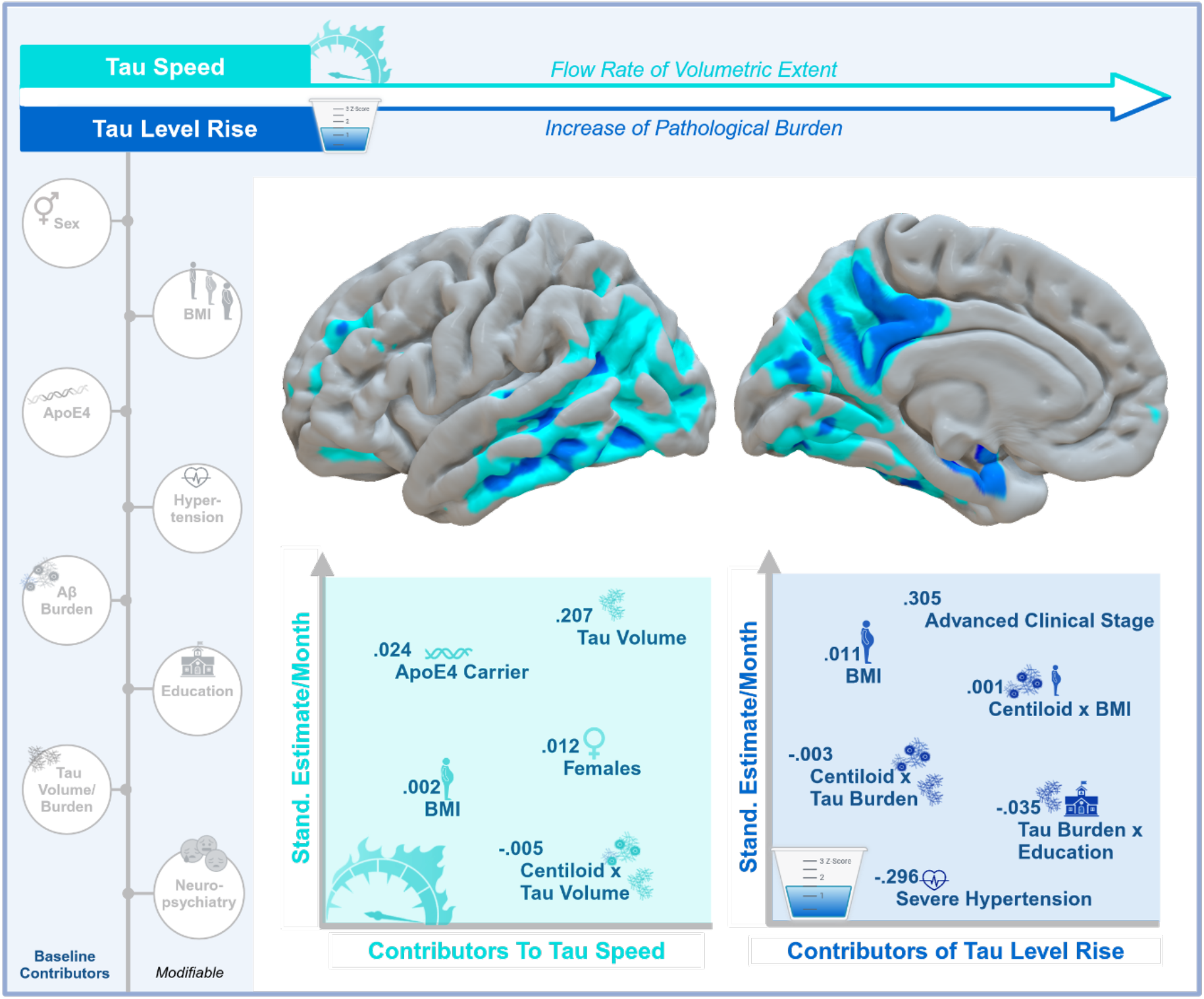
The role of (non-)modifiable factors on tau speed and level rise. Factors that were tested in this study are illustrated on the left. Tau speed (light blue) relates to the volumetric extent of newly affected regions and can be captured as flow rate (volume/month) across time. Tau Level rise (blue) reflects the increase in tau burden in regions that had already been effected by tau pathology at baseline. The brain surfaces depict an example of an male MCI patients, who was 67-year old, had 12 years of education, a BMI of 28.5 and intermediate hypertension at baseline and who became a dementia patient at follow-up. Areas in blue relate to more affected regions and areas in light blue to newly affected regions four years after baseline. The volumetric extent of the light blue areas was used to quantify tau speed, whereas tau level rise was quantified as the change in tau burden in the blue areas. Below the brain surfaces, we have depicted the standardized estimates for the significant fixed and interaction effects relating to tau speed (left) or tau level rise (right).

**Figure 2.**
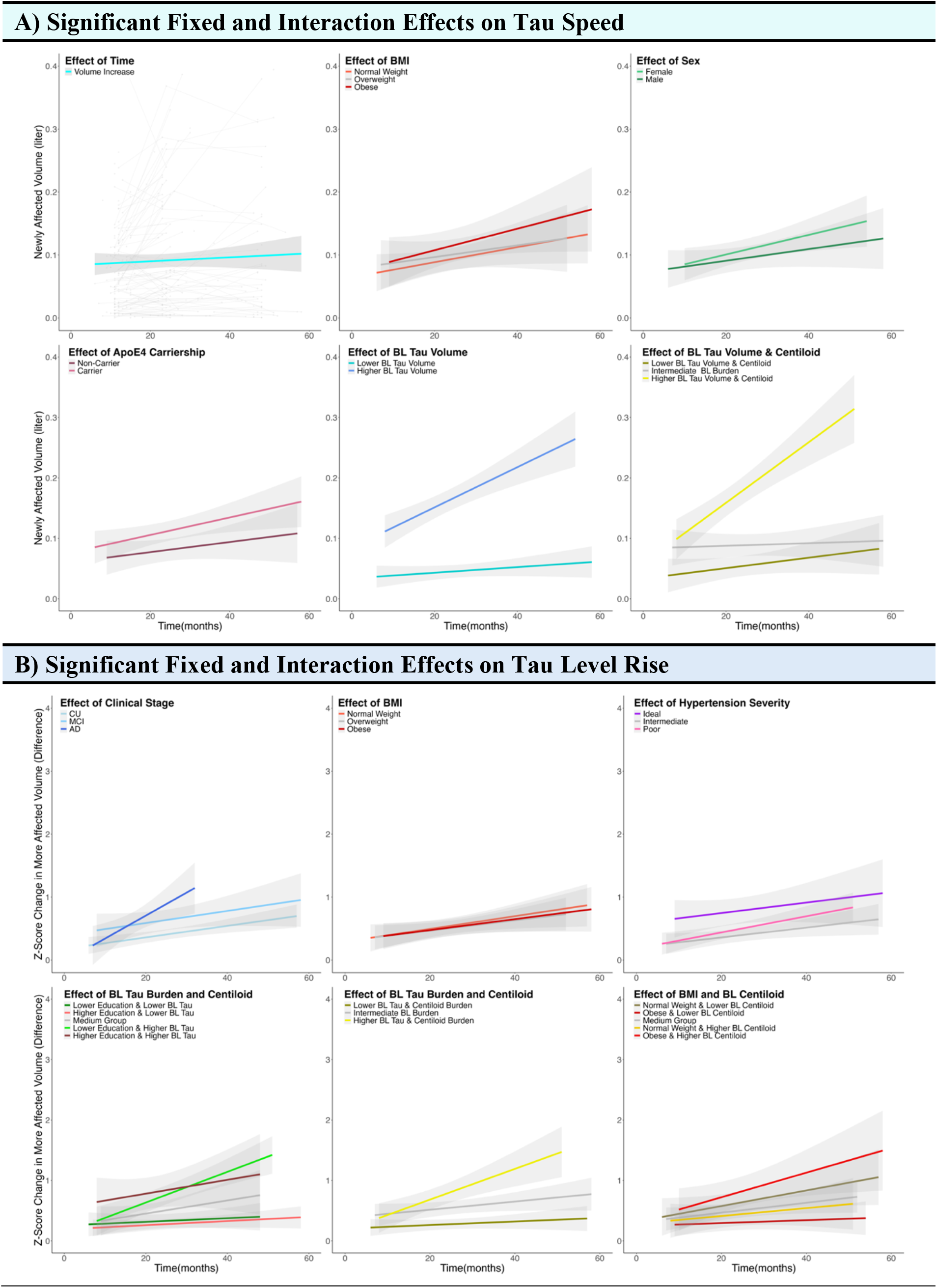
Significant main effects of the variables of interest concerning A) the speed of tau (newly affected volume) and B) tau level rise (increase of tau in more affected regions) are visualized. Continuous variables were converted into grouping variables based on common cut-offs (BMI, education) and the median of the baseline tau volume across all subjects. 95% Confidence intervals are provided for all group effects.

## Discussion

The current study provides novel insights into the intersection of genetic, health risk factors and pathological burden to the spatial progression and regional increase of tau pathology in a cohort of AD-biomarker positive individuals. In line with a previous study ^21^, we showed that baseline amyloid and tau burden affect tau speed and level rise in a dose-dependent manner, with higher initial levels leading to greater progression. Interestingly, we observed that the baseline spatial extent of tau predicted its subsequent volumetric expansion, whereas this relation was not observable for baseline tau burden and its local increase. Potentially, the initial seeding of pathological tau in a certain region causes a self-amplification process, which is independent of its previous seed load ^22^. The greater the volumetric extent of tau, the more regions are affected, and thus the more structural and functional pathways are available, which may facilitate a more rapid tau speed. Predetermined factors, like ApoE4 carriership and female sex, appear to further perpetuate tau’s spatial progression rather than its local increase.

In terms of modifiable risk factors, we found that lower BMI was linked to a slower spatial progression and lower increase in tau burden, whereas greater hypertension severity was linked to a greater increase in tau burden, but not to the spatial expansion of tau pathology.

The mechanistic pathways by which these tested risk factors act on either the progression or increase of tau pathology may partly be explained by impaired waste clearance ^23^. Indeed, poor cardiovascular condition^24^ and a more sedentary lifestyle ^25^ (closely linked to obesity) have both been associated with a worsening of protein clearance from the brain, likely due to a decrease in arterial pulsatility ^26^. Failed clearance, in turn, may lead to progressive aggregation of proteins, such as tau pathology, thereby presumably explaining the effects of hypertension and BMI on tau level rise.

Despite this line of argumentation, we observed a (trend) significant interactive effect of baseline amyloid burden and BMI on tau speed and level rise indicating that the effect of BMI may be driven by its association with amyloid rather than tau burden. Interestingly, it has been shown that obesity in midlife is closely associated with a greater risk of developing AD ^27,28^. Potentially, this risk factor facilitates amyloid-β accumulation during midlife. Indeed, antecedent amyloid burden has been shown to be a key driver of tau pathology aggregation ^29^. It may thus be that the risk of higher BMI on tau spreading is driven by its effects on amyloid aggregation in pre-clinical stages, which subsequently impacts on tau pathology in clinical stages.

In terms of neuropsychiatric symptom severity and education, we only observed baseline associations with pathological burden and an interactive effect of lower education and greater tau burden on tau level rise. This may be due to the nature of these factors as they do not directly interfere with biological pathways, as opposed to BMI or hypertension. Hence, these factors may be more closely related to cognitive function or decline rather than the pathological progress *per se*. Given that education is considered as proxy for cognitive reserve ^30^ (i.e. the preservation of cognition despite increased pathological burden) it may potentially moderate the association between tau speed and level rise and cognitive decline, which remains to be assessed.

A few limitations need to be considered in the interpretation of the current results. First, we only tested interaction effects based on our predefined hypotheses. Thus, we may have missed existing interactive effects across our predictor variables. For instance, given that females suffer from greater cardiovascular risk after menopause^31^ and more severe effects of ApoE4 carriership on tau spreading^32^, future studies may test the association of the tested risk factors stratified by sex. Moreover, in the current study we only focused on four out of the 14 risk factors that have been suggested by the prevention commission^1^. Therefore, it remains unknown whether a single factor carries superior risk for tau progression than others. However, consensus is that the modification of a single risk factor will likely not change the trajectory of AD^1^. Nonetheless, as recently proposed^23^, it will be pivotal to assess the differential contribution of these risk factors to the resistance (i.e. withstanding brain pathology, hence prevention) and resilience (i.e. coping with brain pathology, thus intervention) against AD and its progression.

Overall, this study has provided an initial basis for assessing the contribution of modifiable factors to two aspects of tau pathology, namely its spatial expansion (i.e. tau speed) and its amplification (i.e. tau level rise). The results indicate that education, BMI and hypertension carry differential effects in this regard and closely interact with initial pathological burden. Modification of these factors towards a healthier lifestyle may potentially provide means to slow down and delay disease progression. At what time point in life this modification should take place remains to be tested. Moreover, differential consideration of tau speed and tau level rise may further provide novel and more refined means to study treatment effects of novel drug compounds against AD.

## Acknowledgements

GNB and MH received funding from the Alzheimer Forschungs Initative e.V. In addition, this study was supported by the German Research Foundation (DFG; DR 445/9-1 (AD), CRC1451-C04 Project-ID 431549029 (AD, GNB) & RTG 1960 Project ID 233886668 (VD)) Data collection and sharing for this project was funded by the Alzheimer’s Disease Neuroimaging Initiative (ADNI) (National Institutes of Health Grant U01 AG024904) and DOD ADNI (Department of Defense award number W81XWH-12-2-0012).

## Funding

ADNI is funded by the National Institute on Aging, the National Institute of Biomedical Imaging and Bioengineering, and through generous contributions from the following: AbbVie, Alzheimer’s Association; Alzheimer’s Drug Discovery Foundation; Araclon Biotech; BioClinica, Inc.; Biogen; Bristol-Myers Squibb Company; CereSpir, Inc.; Cogstate; Eisai Inc.; Elan Pharmaceuticals, Inc.; Eli Lilly and Company; EuroImmun; F. Hoffmann-La Roche Ltd and its affiliated company Genentech, Inc.; Fujirebio; GE Healthcare; IXICO Ltd.; Janssen Alzheimer Immunotherapy Research & Development, LLC.; Johnson & Johnson Pharmaceutical Research & Development LLC.; Lumosity; Lundbeck; Merck & Co., Inc.; Meso Scale Diagnostics, LLC.; NeuroRx Research; Neurotrack Technologies; Novartis Pharmaceuticals Corporation; Pfizer Inc.; Piramal Imaging; Servier; Takeda Pharmaceutical Company; and Transition Therapeutics. The Canadian Institutes of Health Research is providing funds to support ADNI clinical sites in Canada. Private sector contributions are facilitated by the Foundation for the National Institutes of Health (www.fnih.org). The grantee organization is the Northern California Institute for Research and Education, and the study is coordinated by the Alzheimer’s Therapeutic Research Institute at the University of Southern California. ADNI data are disseminated by the Laboratory for Neuro Imaging at the University of Southern California.

## Competing Interests

MCH, VD, ED and GNB report no competing interests. TvE reports having received consulting and lecture fees from Lundbeck A/S, Lilly Germany, Shire Germany and research funding from the German Research Foundation (DFG), the Leibniz Association and the EU-joint program for neurodegenerative disease research (JPND). AD reports: Research support by Siemens Healthineers, Life Molecular Imaging, GE Healthcare, AVID Radiopharmaceuticals, Sofie, Eisai, Novartis/AAA, Ariceum Therapeutics; Speaker Honorary/Advisory Boards: Siemens Healthineers, Sanofi, GE Healthcare, Biogen, Novo Nordisk, Invicro, Novartis/AAA, Bayer Vital, Lilly; Stock: Siemens Healthineers, Lantheus Holding, Structured therapeutics, Lilly: Patents: Patent for 18F-JK-PSMA-7 (Patent No.: EP3765097A1; Date of patent: Jan. 20, 2021).

## Supplementary Informations

**S-Table 1.** Correlation results and group comparison of variables of interest. The uncorrected p-value in addition to the FDR-corrected p-value (q) is provided. Green boxes represent significant and orange trend significant associations after FDR correction.

Correlation of baseline factors (n=162):
| Baseline Factors | Centiloid | Tau Burden | Tau Volume | BMI | Education | NPI |
| --- | --- | --- | --- | --- | --- | --- |
| **Centiloid** | 1 |  |  |  |  |  |
| **Tau Burden** | rho= .459<br>p< .001<br>q< .001* | 1 |  |  |  |  |
| **Tau Volume** | rho= .346<br>p< .001<br>q< .001* | rho= .787<br>p< .001<br>q< .001* | 1 |  |  |  |
| **BMI** | rho= -.157<br>p= .024<br>q= .051 | rho= -.080<br>p= .159<br>q= .183 | rho= -.024<br>p= .381<br>q= .408 | 1 |  |  |
| **Education** | rho= -.150<br>p= .029<br>q= .054 | rho= -.115<br>p= .074<br>q= .111 | rho= -.085<br>p= .142<br>q= .183 | rho= .002<br>p= .491<br>q= .491 | 1 |  |
| **NPI** | rho= .219<br>p= .003<br>q= .010* | rho= .209<br>p= .004<br>q= .010* | rho= .208<br>p= .004<br>q= .010* | rho= -.137<br>p= .042<br>q= .070 | rho= -.084<br>p= .147<br>q= .183 | 1 |
| **Hypertension** | H(2; n=162)<br>=.297<br>p=.862<br>q=.862 | H(2; n=162)<br>=3.325<br>p=.190<br>q=.570 | H(2; n=162)<br>=1.742<br>p=.419<br>q=.838 | H(2; n=162)<br>=.354<br>p=.838<br>q=.862 | H(2; n=162)<br>=6.688<br>p=.035<br>q=.210 | H(2; n=162)<br>=.689<br>p=.709<br>q=.862 |
| **ApoE4** | H(1; n=162)<br>=3.628<br>p=.057<br>q=.154 | H(1; n=162)<br>=2.769<br>p=.096<br>q=.154 | H(1; n=162)<br>=4.957<br>p=.026<br>q=.154 | H(1; n=162)<br>=1.563<br>p=.211<br>q=.211 | H(1; n=162)<br>=2.320<br>p=.128<br>q=.154 | H(1; n=162)<br>=2.904<br>p=.121<br>q=.154 |

**S-Table 2.** Results of the LMM for the newly affected volume and the z-score increase in the more affected volume over time. Significant fixed effects are highlighted with an asterisk. The effects of the random slopes are additionally provided for each model. Untandardized beta-coefficients, p-values and confidence intervals (95%) are provided for all effects. The results relate to the subcohort of 90 subjects.

Subcohort- Contributing Effects to Tau Speed and Tau Level Rise
|  | Tau Speed<br>(newly affected volume in liter) |  |  | Tau Level Rise<br>(Z-Score increase in already affected regions) |  |  |
| --- | --- | --- | --- | --- | --- | --- |
| *Fixed effects* | $\beta$ | p | CI | $\beta$ | p | CI |
| **Sex (F/M)** |  |  |  |  |  |  |
| Females vs. Men | .012 | .031* | .001-.023 | .144 | .104 | -.029-.318 |
| **ApoE4 carriership** |  |  |  |  |  |  |
| Carriers vs. Non-carriers | .015 | .011* | .004-.027 | .032 | .721 | -.146-.210 |
| **Clinical Group (CU, MCI, AD)** |  |  |  |  |  |  |
| MCI vs. CU | -.0003 | .955 | -.010-.009 | .076 | .393 | -.100-.254 |
| AD vs. CU | .003 | .633 | -.012-.019 | .321 | .016* | .061-.581 |
| **Hypertension (ideal, intermediate, poor)** |  |  |  |  |  |  |
| Intermediate vs. ideal | -.003 | .728 | -.017-.012 | -.462 | <.001* | -.686- -.239 |
| Poor vs. ideal | .005 | .4928 | -.010-.021 | -.300 | .012* | -.533- -.067 |
| **Age at baseline** |  |  |  |  |  |  |
|  | .0001 | .145 | -.0002-.001 | -.001 | .260 | -.001-.004 |
| **Centiloid** |  |  |  |  |  |  |
|  | .0001 | .256 | -.0002 - .0000 | .001 | .260 | -.001-.004 |
| **Baseline tau volume** |  |  |  | **Baseline tau z-score** |  |  |
|  | .092 | .001* | .040-.145 | -.007 | .883 | -.098-.084 |
| **BMI** |  |  |  |  |  |  |
|  | .001 | .006* | .0003-.002 | .018 | .008* | .005-.030 |
| **Education** |  |  |  |  |  |  |
|  | .001 | .594 | -.002-.003 | -.013 | .448 | -.048-.021 |
| **NPI** |  |  |  |  |  |  |
|  | -.0002 | .776 | -.002-.001 | -.017 | .112 | -.038-.004 |
| **Time (months)** |  |  |  |  |  |  |
|  | .001 | <.001* | .001-.001 | .011 | <.001* | .006-.016 |
| **Random effects** |  |  |  |  |  |  |
| Subject (intercept) | .0002 | .177 | .0000-.0009 |  |  |  |
| Covariance (intercept*slope) | -.0000 | .544 | -.00002-.0000 |  |  |  |
| Time (slope) | .0000 | <.001* | .00001-.0000 | .0002 | .001* | .0001- .0003 |

**S-Table 3.** Interaction effects of the respective models. The fixed effects were the same as in Table 3, but the interaction terms were additionally entered into the respective models. Significant associations are highlighted with an asterisk. The F-statistic and p-values are provided for all interaction effects.

**Subcohort- Interaction of De- and Accelerating Effects on Tau Speed and Tau Level Rise**
| *Interaction effects* |  | F | p |
| --- | --- | --- | --- |
| ***Model 1 - Tau Speed (Newly Affected Volume)*** |  |  |  |
| Centiloid | Tau volume | 14.46 | .050* |
|  | BMI | .727 | .397 |
|  | Education | .048 | .828 |
|  | NPI | .016 | .900 |
|  | Hypertension | .734 | .484 |
| Tau Volume | BMI | .331 | .567 |
|  | Education | .490 | .487 |
|  | NPI | .803 | .374 |
|  | Hypertension | .206 | .814 |
| Hypertension | ApoE4 | .507 | .605 |
| ***Model 2 - Tau Level Rise (Z-Score Change in More Affected Volume)*** |  |  |  |
| Centiloid | Tau volume | 4.37 | .038* |
|  | BMI | 13.35 | <.001* |
|  | Education | 1.23 | .270 |
|  | NPI | .652 | .421 |
|  | Hypertension | .342 | .711 |
| Tau Burden | BMI | 3.19 | .076 |
|  | Education | 3.15 | .078 |
|  | NPI | .217 | .642 |
|  | Hypertension | .426 | .654 |
| Hypertension | ApoE4 | 1.80 | .158 |

